# A Bioluminescent Assay for Quantification of Cellular Glycogen Levels

**DOI:** 10.1101/2024.05.20.595028

**Authors:** Donna Leippe, Rebeca Choy, Gediminas Vidugiris, Hanne Merritt, Kevin T. Mellem, David T. Beattie, Julie C. Ullman, Jolanta Vidugiriene

## Abstract

Glycogen is a large polymer of glucose that functions as an important means of storing energy and maintaining glucose homeostasis. Glycogen synthesis and degradation pathways are highly regulated and their dysregulation can contribute to disease. Glycogen storage diseases are a set of disorders that arise from improper glycogen metabolism. Glycogen storage disease II, known as Pompe disease, is caused by a genetic mutation that leads to increased glycogen storage in cells and tissues, resulting in progressive muscle atrophy and respiratory decline for patients. One approach for treating Pompe disease is to reduce glycogen levels by interfering with the glycogen synthesis pathway through glycogen synthase inhibitors. To facilitate the study of glycogen synthase inhibitors in biological samples, such as cultured cells, a high-throughput approach for measuring cellular glycogen was developed. A bioluminescent glycogen detection assay was automated and used to measure glycogen content in cells grown in 384-well plates. The assay successfully quantified reduced glycogen stores in cells treated with a series of glycogen synthase 1 inhibitors, verifying inhibitor activity in biological systems and validating the utility of the assay for glycogen metabolism studies.

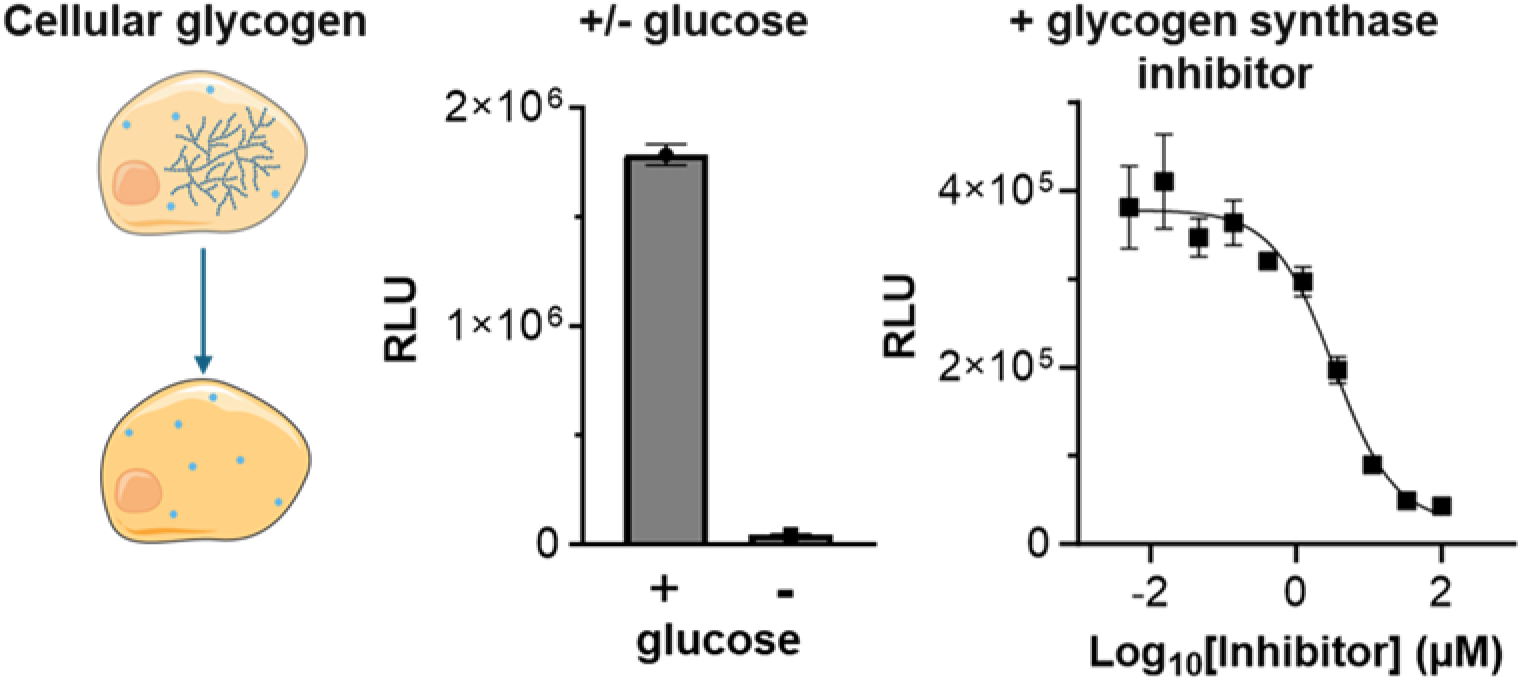

## INTRODUCTION

Glucose is a major energy source for cells and organisms, providing ATP and metabolic intermediates for critical cell processes. Cells store glucose in the form of glycogen, a large intracellular polymer of glucose monomers that can be rapidly mobilized to support energy needs and maintain glucose homeostasis at the cellular, tissue and organismal levels (1,2). Glycogen polymers can contain up to 55,000 glucose units and the processes of glycogen synthesis and degradation are tightly regulated, involving several enzymes (1,2). Metabolic reprogramming in cells can lead to increased glycogen reserves, an adaptation that can potentially sustain cells in-low glucose environments, avoiding the need for external glucose sources. Dysregulation of glycogen metabolism has also been implicated in diseases such as neurodegenerative diseases (3), cancer (4,5), diabetes (6) and glycogen storage diseases (GSD’s; 7). Glycogen’s important role in health and disease makes it a target of interest for various therapies (8).

There are several types of glycogen storage diseases with different enzyme defects and clinical presentations. Pompe disease is glycogen storage disease II, and it is characterized by mutations in the acid alpha-glucosidase enzyme (GAA) that alter its ability to degrade glycogen (9,10). Glycogen degradation occurs through two main pathways: cytosolic and lysosomal (1). GAA is responsible for lysosomal degradation and with reduced or non-existent GAA activity, glycogen builds up in the tissues of patients with Pompe disease, leading to severe musculoskeletal symptoms and shortened lifespan (9,10).

Substrate reduction therapy (SRT) has been proposed as a novel therapeutic hypothesis to test for the treatment of patients with Pompe disease (11). This approach targets inhibition of Glycogen Synthase 1 (GYS1) to reduce glycogen in the muscle tissues affected in Pompe disease (12). GYS1 is the rate limiting enzyme in glycogen synthesis within muscle cells and inhibition of GYS1 has been demonstrated to reduce glycogen accumulation in muscles cells (12). Two groups have published reports describing the synthesis and characterization of small molecule inhibitors of GYS1 activity (12, 13) and both used initial screening platforms which relied on in vitro reactions with recombinant enzymes. While in vitro biochemical enzyme assays are useful for deriving IC_50_ values they are unable to predict cell permeability or cell stability.

Many commonly used cellular glycogen assays rely on colorimetric readouts with limited dynamic range and poor sensitivity which necessitates large sample input. These features hinder the application of these methods for cell-based glycogen measurements in high-throughput formats. To address this need in the research community, we evaluated a bioluminescent glycogen assay with a broad assay range and greater sensitivity for compatibility with high-throughput screening protocols. The bioluminescent glycogen assay is based on converting glycogen to glucose and utilizing a bioluminescent NAD(P)H detection method to measure the released glucose. The bioluminescent NAD(P)H technology (14) has previously been applied for the sensitive detection of various cellular metabolites and is compatible with automated systems and high-throughput screening processes (15, 16).

In this paper we describe the development of a high-throughput method for measuring glycogen to enable the characterization of compounds for their ability to inhibit glycogen synthesis in cell-based systems. Multiple compounds were tested during development, including MZ-101 which was recently published in a study demonstrating its ability to selectively inhibit GYS1, reducing accumulated glycogen in a mouse model of Pompe, and demonstrating the potential therapeutic efficacy of SRT for the treatment of Pompe disease (12). Our results validate the bioluminescent method as a robust assay to quantify changes in cellular glycogen levels supporting the utility of this assay both as a screening paradigm to identify modulators of glycogen levels and for studying basic biology of glycogen metabolism in living cells.

## RESULTS AND DISCUSSION

### Glycogen Assay Principle and Performance

The bioluminescent glycogen assay quantitates cellular glycogen after cell lysis and subsequent enzyme conversion of glycogen to glucose. The assay uses glucoamylase to enzymatically digest glycogen into glucose monomers followed by glucose measurement (Fig 1A). Glucose measurement is achieved through coupled-enzyme reactions using a bioluminescent system for NADH detection that includes glucose dehydrogenase and NAD, providing a sensitive and specific measurement of glycogen-derived glucose (Fig 1B). The result is a light signal directly proportional to the amount of glycogen in the sample. Both the glucose dehydrogenase and NADH detection reactions occur simultaneously within a single “glucose detection reagent” allowing for glucose measurement in a single step.

**Figure 1.**
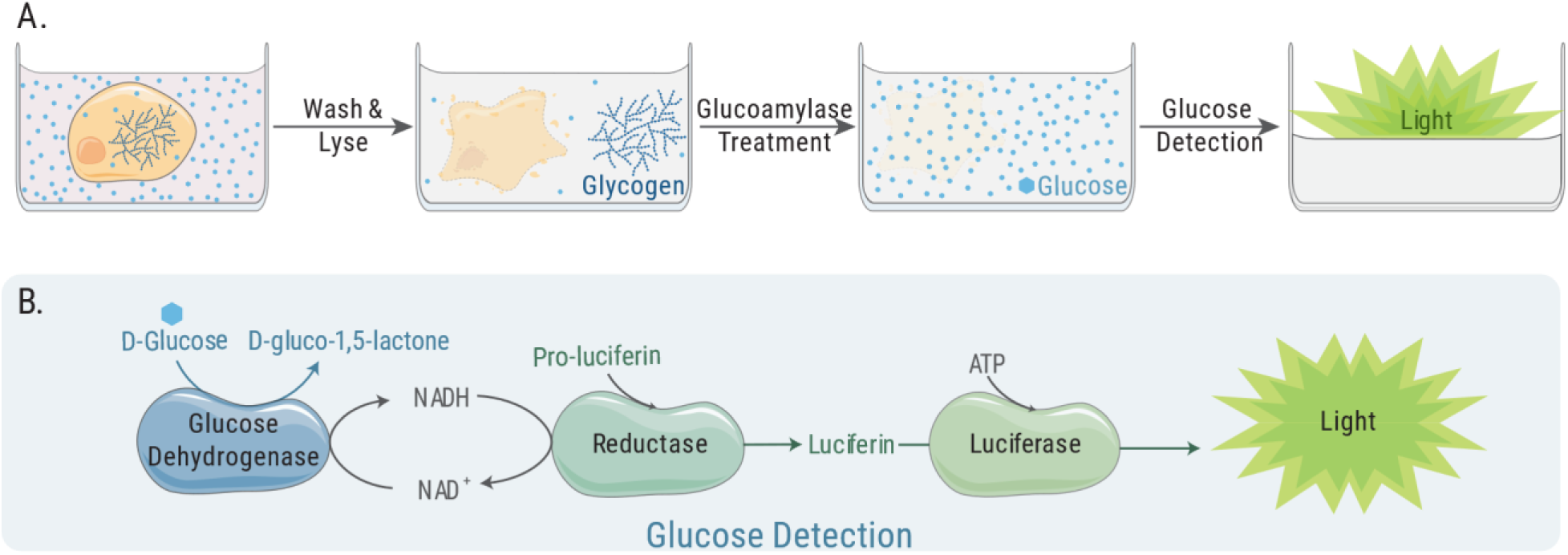
Schematic of the process for glycogen detection using the bioluminescent method. (A) Cells are washed and lysed before glycogen is digested into glucose monomers for subsequent glucose detection. (B) Bioluminescent assay for detecting released glucose.

When measuring glycogen in cells and other biological samples the presence of basal glucose must be considered. The addition of glucoamylase is a key step for distinguishing the signal originating from glycogen and the signal originating from basal glucose levels in cells and tissues. To determine the amount of glycogen in samples that also contain glucose, two parallel reactions are performed: one with glucoamylase and one without glucoamylase. The glycogen-specific signal is obtained by subtracting the “no glucoamylase” reaction signal from the signal of the glucoamylase-treated reaction.

In addition to basal glucose levels, commonly used cell culture media also contains high levels of glucose that will contribute to the assay signal, requiring washing steps before conducting any measurements. Once the excess glucose has been removed, the cells can be lysed (Fig 1A). We use strong acidic conditions to lyse cells. This lysis method also achieves minimal assay background by aiding in (1) cellular enzyme inactivation, including endogenous dehydrogenases, which might produce NADH from the NAD added to the glucose detection reagent and (2) cellular NADH and NADPH degradation. Though endogenous levels of NADH and NADPH are low, their removal ensures lowest assay background and greatest sensitivity.

Key parameters of the glycogen assay were determined using a titration of purified glycogen. The assay was linear over a 100-fold range of 0.02 to 20 µg/ml (Fig 2A). There was no signal above background in the absence of glucoamylase. The assay had a signal-to-background of 1.4 and a signal-to-noise ratio of 18 at the lowest concentration tested (0.02 µg/mL), which confirmed its high sensitivity (Fig S1). Based on reported values (17), this was calculated to be a suitable range for measuring glycogen in cells cultured in 96– and 384-well plates.

**Figure 2.**
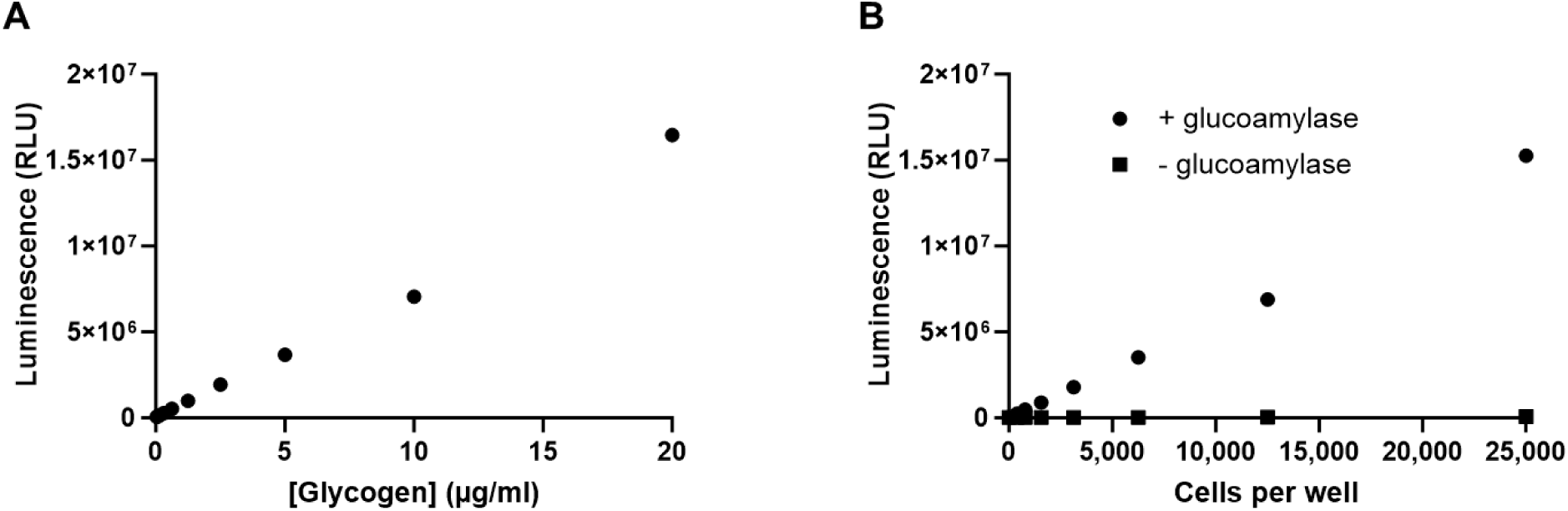
Detection of glycogen. (A) Titration of a purified stock of glycogen. Two-fold serial dilutions of glycogen were prepared in PBS. Aliquots (25 µl) of each dilution were transferred to quadruplicate wells of a 96-well plate and assayed as described in the Methods section. Luminescence was recorded as relative light units (RLU). The average RLU are plotted. Error bars are +/− 1 s.d. Percent CVs were ≤5%. (B) Titration of HeLa cells for glycogen measurement in the presence and absence of glucoamylase. HeLa cells were cultured in complete growth medium overnight before washing and cell lysis. The cell lysates were diluted two-fold and the lysed cell equivalents per well is plotted on the x-axis. Each data point represents the average of three replicates. Error bars are +/− 1 s.d.

### Measurement and Modulation of Glycogen Levels in HeLa Cells

In this study, we aimed to develop a high-throughput method for glycogen detection in mammalian cells. Glycogen levels vary across different cell types (17) and are influenced by media composition, especially glucose availability. HeLa cells are a very commonly used adherent cell line for biological research and screening tool for drug development; it is for these reasons we chose HeLa cells as the model system to develop the assay. To measure the range of glycogen that could be detected in HeLa cells, cells were grown in complete DMEM growth medium containing 25 mM glucose and subsequently washed in PBS, counted, and lysed as described in the Methods section. After lysis, the cell lysate was serially diluted by two-fold, and 25 µl of each dilution was added (per well) to a 96-well assay plate for glycogen measurement. A glycogen standard curve was included on the same assay plate for glycogen quantification.

The glycogen assay detected cellular glycogen present in as few as 300 cells and up to 25,000 cells in a linear manner (Fig 2B). Parallel reactions lacking glucoamylase were performed to detect cellular glucose. As observed previously with other cancer cell lines (15), the glucose levels were not above assay background (Fig 2B) presumably due to its rapid conversion to glucose 6-phosphate in the cell. Given that the basal glucose concentrations were below the detectable range, the glycogen content in the cell lysates could be calculated directly from the readings of the glucoamylase-treated samples. The amount of glycogen in 25 µl lysate containing 25,000 cells was calculated to average 19 µg/ml, corresponding to approximately 19 pg/cell.

Our goal was to develop a sensitive assay with a large dynamic range that could be used to measure glycogen levels in cells grown in 384-well plates with the aim of applying this method as a screening tool to support identification and characterization of small molecule modulators of glycogen synthesis and breakdown. To optimize the assay window, we analyzed how HeLa cell glycogen levels change in response to varied glucose concentrations in culture media. HeLa cells were incubated overnight in flasks with culture medium containing either 25 mM glucose, the same concentration in complete DMEM growth medium, or starvation medium containing 0 mM glucose; cells were then harvested and processed for the assay. As shown in Figure 3A, glycogen was dramatically depleted in cells cultured without glucose. Glycogen levels for 25,000 cells dropped 112-fold from 19 µg/ml to 0.17 µg/ml (Fig 3A). The 0.17 µg/ml concentration was still detectable above assay background which illustrates the importance of assay sensitivity to observe the full range of glycogen levels and not limit signal windows.

**Figure 3.**
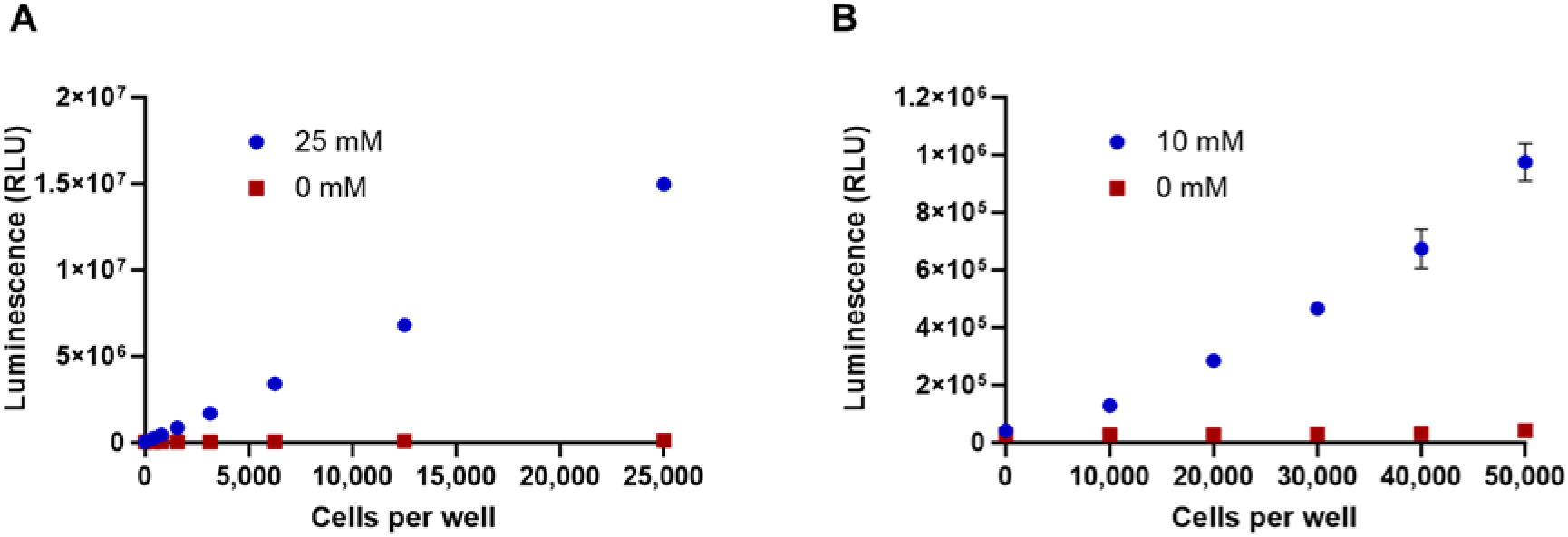
Modulation of glycogen in HeLa cells. (A) HeLa cells were cultured overnight in flasks in either medium with 25 mM glucose or medium lacking glucose (0 mM glucose) and then assayed for glycogen. The number of lysed cell equivalents assayed per well of a 96-well plate is indicated on the x-axis. (B) After an overnight starvation in flasks, HeLa cells were collected and washed and plated in the wells of a 96-well plate at the indicated cell number per well. They were then incubated overnight in the presence of medium containing 10 mM or 0 mM glucose before the glycogen assay. In both panels each data point represents the average of quadruplicate wells. Error bars are +/− 1 s.d.

When glycogen depleted cells were reintroduced to medium containing glucose, glycogen reaccumulated (Fig 3B). Reaccumulated glycogen levels were dependent on the glucose concentration in the culture medium (Fig S2). Considering that glucose in medium needs to be removed by washing with PBS before glycogen measurements, we chose a concentration (10 mM) that could be removed with the least washing and still result in high glycogen accumulation for future experiments. Though glycogen was restored in cells during the overnight incubation, it did not reach the levels previously observed in cells grown in complete growth medium that had not been starved (Fig 2B). Wells with 30,000 cells contained 0.59 µg/ml glycogen, ∼32-fold less than observed with cells cultured in complete medium. However, despite the lower levels, good signal windows were maintained. The ratio of signals from the 10 mM and 0 mM glucose samples was 23, 16 and 11 with 50,000 cells, 30,000 cells and 20,000 cells per well, respectively.

### Development of Automated Protocol for Glycogen Detection

Automation and miniaturization of the bioluminescent glycogen assay are key for high-throughput applications, such as compound screening. Accurate detection of intracellular glycogen relies on the enzymatic conversion of glycogen into glucose and the distinction between glucose originating from glycogen versus basal and media glucose. Therefore, medium removal and washing steps are important. For instance, when cells are cultured in medium containing 10 mM glucose, the glucose concentration must be reduced by approximately 10,000-fold to ≤1 µM to have accurate intracellular glycogen measurement. Incomplete glucose removal can lead to high background and well-to-well variability. In addition, this reduction in glucose concentration by successive washing steps must not disrupt the cell monolayer which would result in cell loss.

To optimize glucose removal and washing steps, HeLa cells were dispensed into 384-well plates at densities of 15,000 and 7,500 cells per well using a Multidrop ™ Combi nL Reagent dispenser and allowed to adhere overnight to ensure a stable cell monolayer formation. The following day, medium was removed, and cells were washed with PBS using a Tecan Freedom EVO^®^ liquid handling system equipped with a MultiChannel Arm™ (MCA) 384. The washing steps were optimized by evaluating different tip height and aspiration/dispensing speeds. The efficiency of glucose elimination from the wells was assessed by measuring glucose levels in the washing solution after completing the cell wash cycles. Furthermore, to evaluate the impact of the washing protocol on cell viability and adherence, the CellTiter-Glo^®^ Luminescent Cell Viability Assay (CellTiter-Glo^®^ Assay) was employed post-wash. The results indicated that performing 8 wash cycles with the liquid handling tips set at a height of approximately 1.5-2 mm from the bottom of the well, combined with an aspiration and dispensing speed of 20 µl/sec, achieved the most efficient glucose removal with minimal effect on cell monolayer (Fig S3).

The performance of the automated assay was further validated by measuring glycogen in HeLa cells plated at 20,000 cells per well in 384-well plates. To monitor the well-to-well variability, 24 wells at different plate positions were used. The cells were plated, incubated overnight, washed, and lysed. Glucoamylase was added and after digestion, a small volume (8 µl) was transferred to a 384-well low volume (LV) plate for the glycogen assay. An aliquot was also transferred to a second 384-well LV plate for viability readings. Residual glucose from media and basal intracellular glucose (previously shown to be not detectable) were measured in a parallel reaction without glucoamylase.

As shown in Figure 4, the optimized washing protocol efficiently reduced extracellular glucose to background levels (Fig 4B). A 100-fold increase in signal was measured in glucoamylase-treated samples with low well-to-well variability (∼9%; Fig 4A), while maintaining the cell monolayer and viability (Fig 4C).

**Figure 4.**
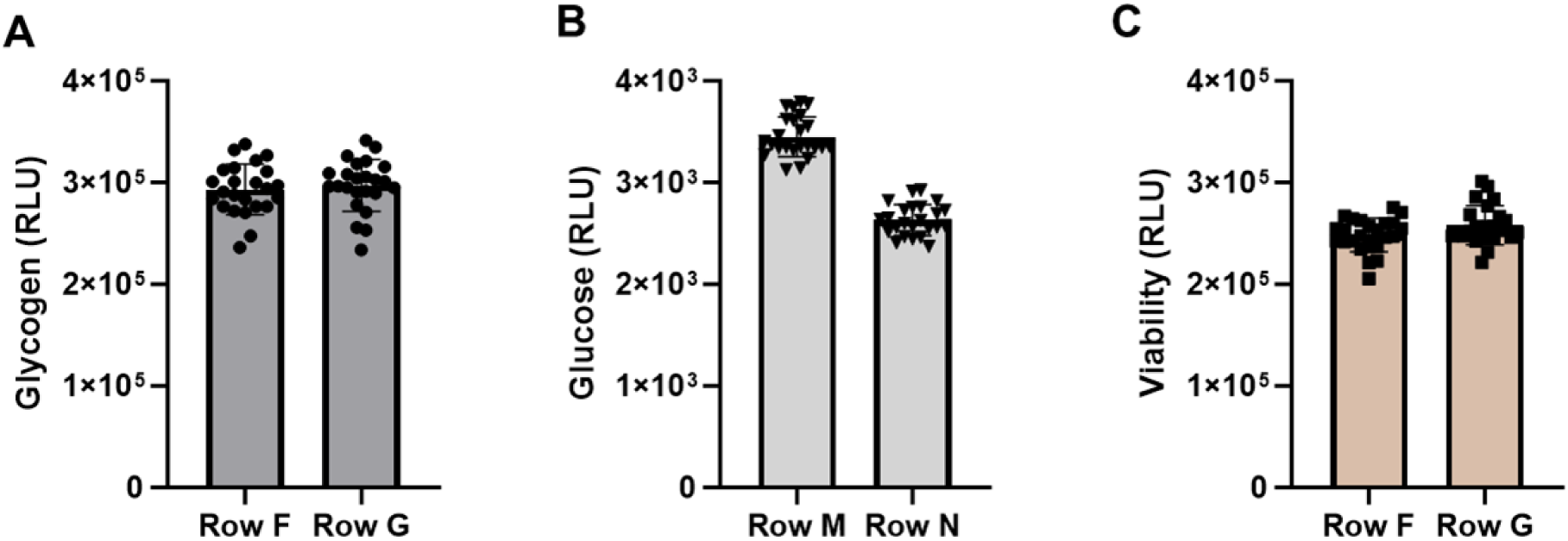
Performance of the automated protocol. The steps in the method were validated using HeLa cells plated in 384-well plates in medium containing 10 mM glucose. The cell density was 20,000 cells per well. Each row contained 24 wells. After washing and cell lysis several parameters were checked. (A) Glycogen measurement in cell lysates. (B) Measurement of basal and residual glucose. For glucose measurements the glucoamylase enzyme was not included. (C) ATP measurement in cell lysates using the CellTiter-Glo^®^ Assay.

### Assay Optimization to Evaluate GYS1 Inhibitors

To support quantification of IC_50_ values for compounds designed to inhibit newly synthesized glycogen, the growth conditions of the HeLa cells were adapted to provide a robust assay window. Briefly, this involved growing the cells in a glucose-free medium to first deplete glycogen stores followed by replenishment of glucose media with or without small molecule inhibitors of GYS1. This approach was not only well tolerated by the cells but importantly it led to a robust increase in the dynamic range of the assay. To further streamline the high-throughput screening workflow and avoid an in-plate wash step, the cells were cultured in glucose-free conditions overnight in flasks and were washed in bulk before dispensing into 384-well plates in media with 10 mM or 0 mM glucose. The final conditions were determined by evaluating the dynamic range of glycogen as a function of cell density (Fig 5A) and cell viability (Fig 5B). Glycogen data for HeLa cells plated at 3 densities, 20,000, 15,000 and 7,500 cells per well, are presented in Figure 5. While 20,000 cells per well had the highest amount of glycogen per well, the level in 20,000 starved cells was also higher, narrowing the signal window (Fig 5C). Therefore, a density of 15,000 cells per well was selected for further studies. A viability assay, the RealTime-Glo™ MT Cell Viability Assay (RealTime-Glo™ Assay; 18) was incorporated into the final protocol to serve as an indicator of both cell health and well-to-well cell plating consistency (Fig 5B).

**Figure 5.**
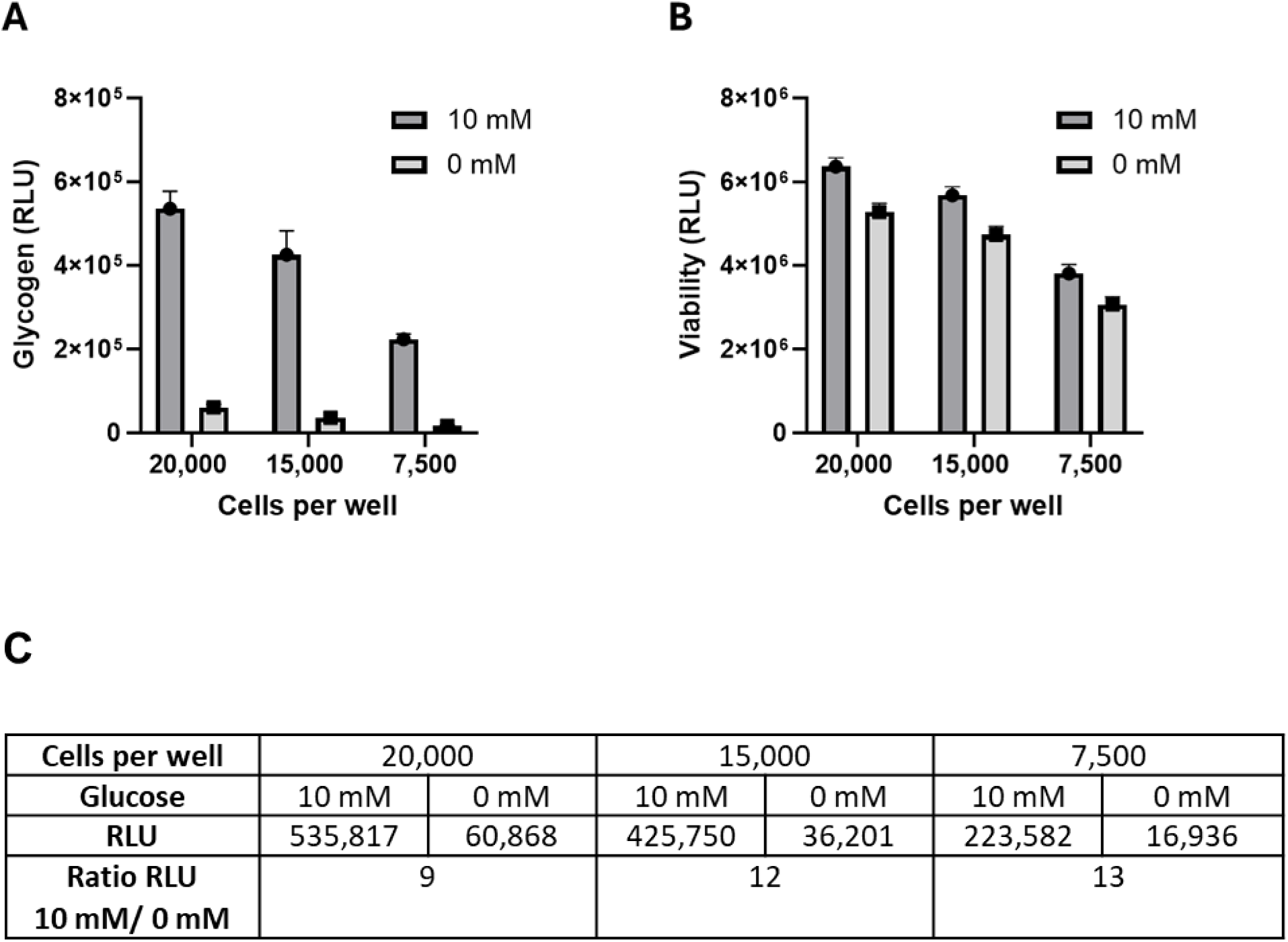
Optimization of cell density. Multiple cell densities were tested to optimize the signal window between cells grown in the presence of 10 mM glucose and those grown in the absence of glucose (0 mM). Cell densities were 20,000, 15,000 and 7,500 cells per well, in 384-well plates (40 wells each). (A) Measurement of glycogen for three cell densities in two media. (B) Cell viability after incubation but prior to cell washing and lysing using the RealTime-Glo™ Assay. (C) Average RLU values and the ratio of RLU values from cells grown in 10 mM and 0 mM glucose. The average RLU are plotted. Error bars = +/− 1 s.d.

### Assay Validation with Small Molecule Inhibitors of GYS1

Recently Ullman, et al. (11) reported on the synthesis and preclinical characterization of the small molecule MZ-101, a potent and selective inhibitor of GYS1 in vitro and in vivo. To validate the glycogen assay for high-throughput applications, we applied the optimized protocol (Table S1) using three compounds that were also shown to inhibit recombinant GYS1 in a biochemical assay (data not shown; 19). The three compounds (referred to as model compounds C1, C2 and C3) were tested in dose-response curves using serial two-fold dilutions starting at 100 µM (final concentration) with a final concentration of 1% DMSO. The compounds were incubated with the cells during the glucose repletion step for 24 hours before cells were lysed and glycogen levels measured. To evaluate intra– and inter-plate variability, each compound was tested in two different plates in two positions on each plate for a total of 4 experimental replications per compound (Fig 6A). Data from the assay were used to derive IC_50_ values for all three compounds: 3.1 to 4.7 µM, 4.5 to 7.8 µM and 10.6 to 12.4 µM for compounds C1, C3 and C2, respectively (Fig 6B). The RealTime-Glo™ Assay results verified that the decrease in glycogen levels was not attributable to compound toxicity or reduced cell growth and thus was due to inhibition of glycogen synthesis. A decline in cell viability was observed only at the highest 100 µM concentration, with viability percentages of 77%, 65% and 75% obtained with compounds C1, C2 and C3, respectively (Fig 6A).

**Figure 6.**
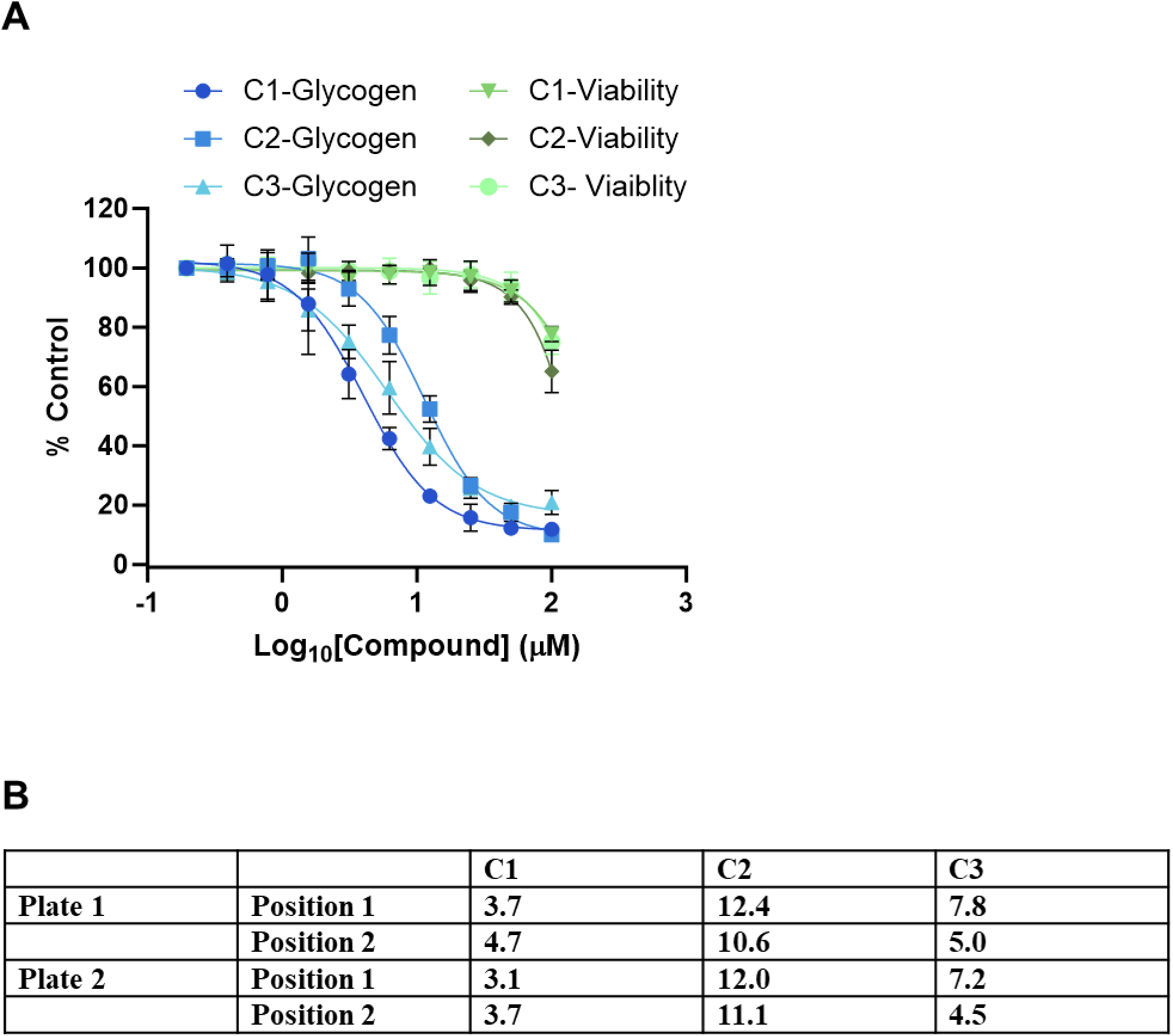
Control compound effects on glycogen levels. Two-fold serial dilutions of each of the three compounds were made in in medium starting at 100 µM (final concentration in well with cells). A total of 10 concentrations were tested in quadruplicate. Each of the three control compounds was tested in two positions on Plate 1 and two positions on Plate 2. (A) Percent glycogen inhibition and percent viability for each inhibitor. The four sets of data for each inhibitor were averaged. The results are expressed as percent of the control cells incubated in glucose in the absence of inhibitor. (B) IC_50_ values (µM) calculated from the 4 individual sets of data for each inhibitor.

### Validation of the Assay for High-Throughput Screening of GYS1 Inhibitors

To further validate the method by determining if the assay could distinguish compounds of different potencies, we expanded our study and screened 20 additional compounds found to have different activities in the biochemical assay, including MZ-101 (12, 19). Four 384-well plates were used for the screen. Each plate contained 5 test compounds and included reference compound C1 as a quality control. The layouts of the compound dilution 96-well plates and corresponding 384-well assay plates are shown in Supplemental Information Figure S4. The screening protocol is outlined in Table S1. All compounds were active and inhibited glycogen synthesis in a dose-dependent manner in cells (Fig 7A). The compounds exhibited different potencies with IC_50_ values ranging from 0.07 to 4.6 µM (Fig 7B). The signal window for each compound was calculated as a ratio of light units at high and low compound concentrations. All compounds had signal windows ≥5 and the ratios ranged from 6.8 to 13.8 (Fig 7B). The test compounds had only slight effect on cell viability, with <20% cell death at the highest concentration, except for 1 of the 20 compounds which had >60% cell death at the highest concentration (data not shown). Each assay plate incorporated quality controls which were used to qualitatively evaluate the overall performance of the assay including: a titration of control compound C1, vehicle (DMSO)-treated cells grown in starvation medium, and vehicle-treated cells grown in glucose-containing medium (Fig S5). The IC_50_ values for compound C1 across the four test plates were 3.1, 3.2, 3.1 and 3.5 µM, which aligned with the IC_50_ values obtained during the small-scale screening tests (Fig S5).

**Figure 7.**
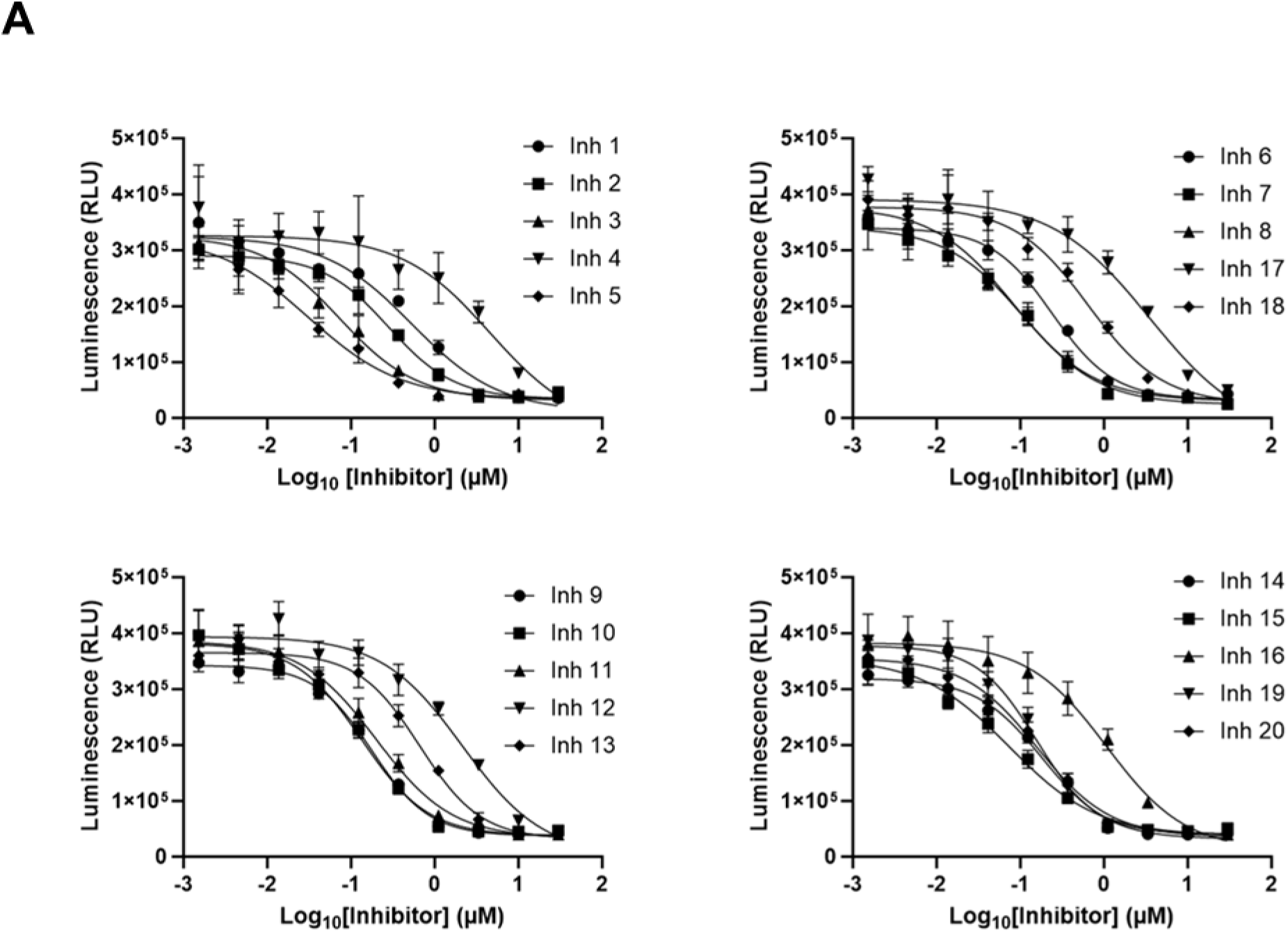
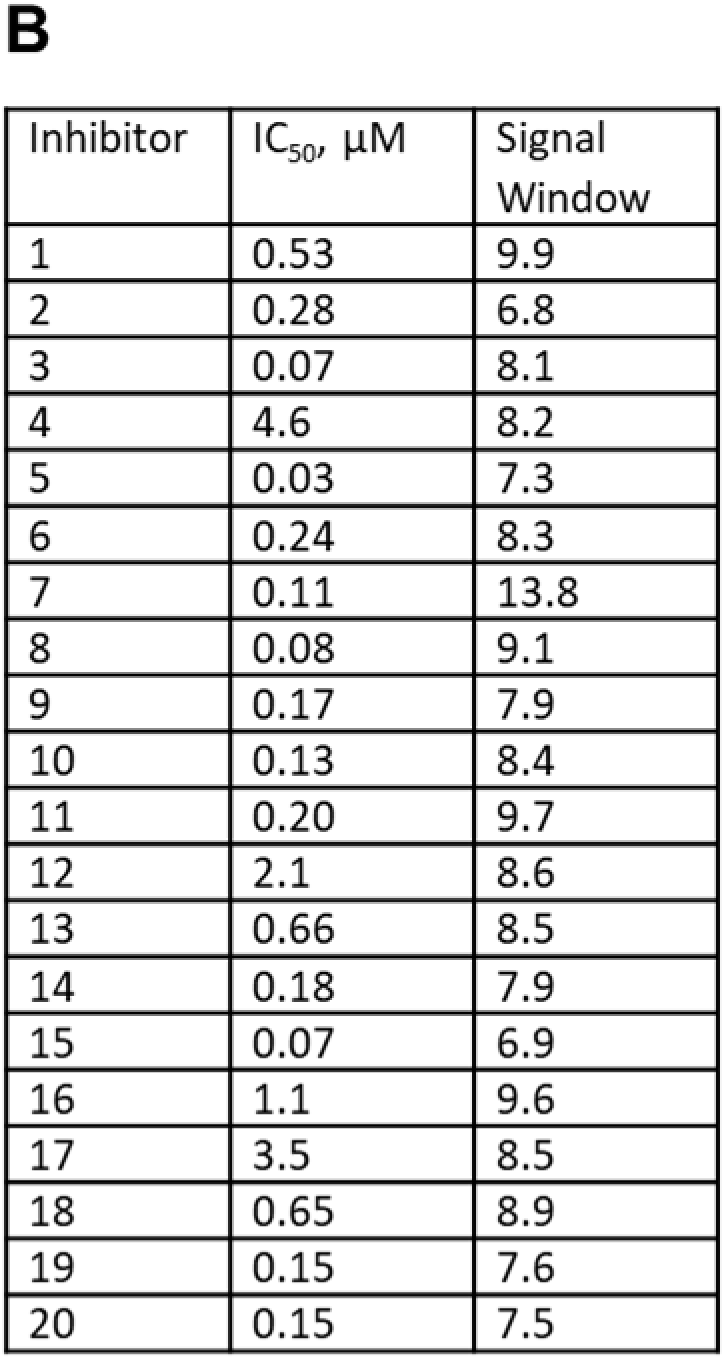
Dose response curves for 20 GYS1 inhibitors and their IC_50_ values. Three-fold serial dilutions of each of the 20 compounds were made in medium starting at 30 µM (final concentration in well with cells). A total of 10 concentrations were tested in quadruplicate. Five inhibitors were tested per plate, for a total of 4 plates. (A) Dose response curves for each inhibitor. The average RLU are plotted. Error bars are +/− 1 s.d. (B) IC_50_ values and signal windows calculated for each of the inhibitors.

The assay’s robustness was further validated by a calculated Z’ factor of 0.74 (Fig 8; 20). This value was derived from comparing luminescence between starved cells and those returned to glucose-containing medium across all four plates. The Z’ values for the four screening plates were 0.81, 0.79, 0.79, and 0.78 for plates 1 through 4, respectively, demonstrating excellent assay quality and reproducibility. Also, the average RLU value for starved cells was above the assay background (35,519 +/− 6,407 as compared to 11,111 +/− 1,788) an important feature for inhibitor screening necessary to observe maximum inhibition. The average RLU value for cells in 10 mM glucose was 350,201 +/− 21,184, resulting in an assay window of 9.9-fold. The 20-compound screen showcases the assay’s ability to discern compounds with IC_50_ values ranging from nanomolar to micromolar (30 nM to 12.4 µM) values. The assay shows robustness across multiple plates while maintaining high reproducibility and consistency in IC_50_ determinations.

**Figure 8.**
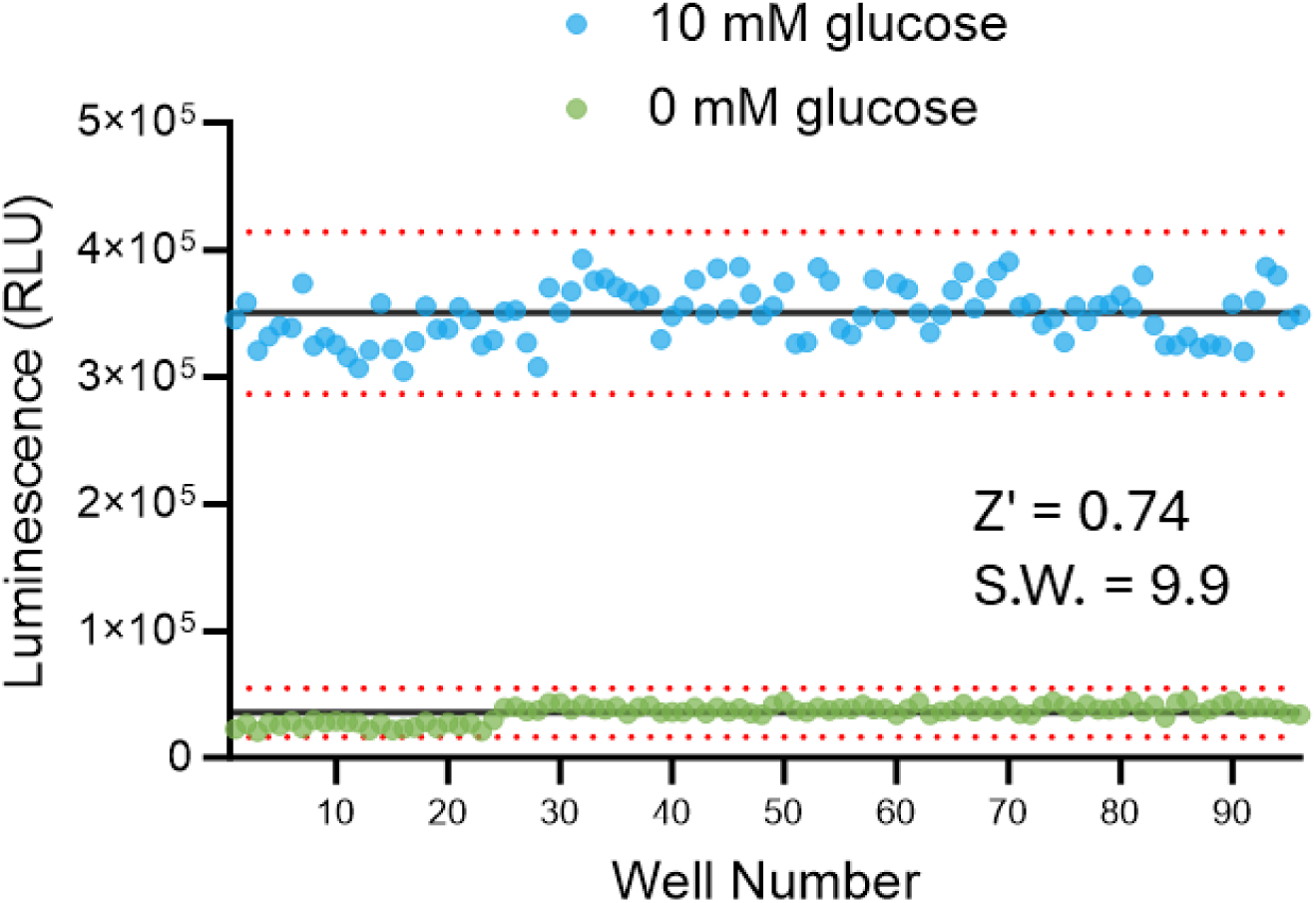
Z′ factor for the automated glycogen detection assay. Light signals from control wells containing cells plated in 10 mM glucose or 0 mM glucose are plotted. Each of the four assay plates had 24 wells of each control for a total of 96 wells (well number). The data were used to calculate the Z′ value and the signal window (S.W.; ratio of 10 mM /0 mM average RLU). The dotted lines are +/− 3 s.d. for each data set.

## CONCLUSIONS

New therapies for rare and common metabolic diseases have led to a resurgence of interest in technologies to monitor glycogen metabolism. The development and validation of the novel bioluminescent assay reported herein highlights a new tool for the study of glycogen metabolism and its dysregulation in disease states. The assay was successfully applied to the characterization of GYS1 inhibitors, including MZ-101 (12), and supports more efficient selection and development of potential SRTs for the treatment of patients with Pompe disease. Our results demonstrate the assay’s sensitivity and specificity in quantifying cellular glycogen levels across a wide dynamic range as well as its compatibility with high-throughput screening, thus making it a valuable tool for both basic science research as well as drug discovery.

## METHODS

### Cell Culture

HeLa cells (ATCC Cat. #CCL-2) were stored in liquid nitrogen. For experiments, cells were thawed and used at passage number ≤15 post-thaw. Cells were propagated in 15 ml complete growth medium consisting of DMEM (Gibco Cat. #11995-065) and 10% FBS (VWR Cat. #89510-194) in T75 flasks. For growth at defined glucose concentrations, DMEM minus glucose, glutamine, pyruvate, and phenol red (Gibco Cat. #A14430-01) supplemented with 10% FBS and 1X GlutaMax™ Supplement (Gibco Cat. #35050-061) was used as a base medium. Glucose (diluted from Gibco Cat. #A24940-01) was added to the base medium to achieve the concentrations indicated in each experiment. For starvation medium, no glucose was added (0 mM glucose). For recovery medium, 20 mM glucose was added, resulting in 10 mM glucose final concentration when added to cells at a 1:1 ratio (v/v). All media used for compound dilutions contained 20 mM glucose and 2% DMSO (Sigma Cat. #D2650-5×5ml).

### Glycogen Detection Assay

Glycogen was measured using the Glycogen-Glo™ Assay (Promega Cat. #J5051) following the manufacturer’s instructions. Briefly, before measuring cellular glycogen, medium was removed, and cells were washed with PBS (Dulbecco’s Phosphate Buffered Saline; Gibco Cat. # 14190-144) to remove residual glucose from the medium. Cells were then lysed by adding 0.3N HCl followed by the addition of 450 mM Tris pH 8.0. The recommended volume ratio of cells in PBS to acid and base is 1: 0.5: 0.5. Glucoamylase solution was prepared by diluting glucoamylase enzyme in glucoamylase buffer (100 mM sodium acetate, pH 5.2), and added to the cell lysate at a 1:1 volume ratio (e.g., 25 µl glucoamylase solution added to 25 µl cell lysate in a 96-well plate or 30 µl added to 30 µl cell lysate in a 384-well plate). The digestion reaction continued for 1 h at room temperature at which time an equal volume of glucose detection reagent (containing glucose dehydrogenase, NAD and the components of the bioluminescent NAD(P)H detection technology) was added. Luminescence was recorded after 90 min at room temperature. For the automated cell-based experiments presented in this paper, two modifications were made: the cell lysates for the automated protocol were frozen at –20°C overnight before the glucoamylase was added and (2) the glucoamylase reaction was incubated for 1 h at 37°C.

### Automated Protocol

For the automated protocol, cells were plated in 384-well tissue culture-treated microplates (Corning Cat. #3707, clear bottom, white-walled). Cells were dispensed into 384-well plates using a MultiDrop™ Combi nL Reagent Dispenser (ThermoFisher Scientific). Cell culture medium was removed and cells were washed with PBS using a Tecan Freedom EVO^®^ Robotic Platform equipped with the MultiChannel Arm™ (MCA) 384. The MultiChannel Arm™ (MCA) 384 was used to add HCl and Tris, and transfer the 8 µl aliquots of cell lysates. The glucoamylase solution, glucoamylase buffer, glucose detection reagent and viability reagents were dispensed using the Multidrop™ Combi nL Reagent Dispenser. Bioluminescent assays were performed in 384-well LV plates (Corning #4512, white, opaque plates). Luminescence was recorded using a Tecan infinite M200 PRO Multiplate Reader.

To optimize the automated protocol, cells were dispensed at the cell densities indicated for each experiment and incubated overnight in 50 µl recovery or starvation medium. The next day media was removed and the cells were washed with 70 µl PBS per wash cycle. At the end of the washing protocol, ∼15 µl PBS remained in the wells. Acid, 7.5 µl 0.3N HCl, was added per well, followed by addition of 7.5 µl 450 mM Tris pH 8.0. Plates with cell lysates were sealed and stored overnight at –20°C. The following day, 30 µl glucoamylase solution was added per well. After digestion, 8 µl was transferred to a 384-well LV plate and 8 µl of glucose detection reagent was added for glucose measurements.

The effectiveness of the washing protocol was assessed by monitoring glucose levels in parallel wells. In this case 30 µl of glucoamylase buffer without glucoamylase was added to the cells. A sample (8 µl) was transferred to a 384-well LV plate for glucose measurement. A cell viability assay was included to confirm that cells were not dislodged and lost during the washing protocol. An additional 8 µl of cell lysate was transferred to a 384-well LV plate and ATP was measured by adding 8 µl CellTiter-Glo^®^ Luminescent Cell Viability Assay (Promega Cat. #G7571).

### Screening Protocol

Cells were propagated overnight in starvation medium in T75 flasks. Cells were collected (using trypsin/EDTA), washed with PBS and dispensed in starvation media at 15,000 cells in 25 µl per well of a 384-well plate. All inhibitor compounds were from Maze Therapeutics (South San Francisco, CA) and were supplied as 10 mM stocks in DMSO. They were each assayed in 10-point dose response curves. The three control compounds were diluted to 200 µM and serially diluted two-fold. The twenty additional compounds were diluted to 60 µM and serially diluted three-fold. All compound dilutions were in recovery medium containing 20 mM glucose and 2% DMSO. Each dilution (25 µl) was added to quadruplicate wells of the 384-well plate containing dispensed cells. Cells were incubated for 24 h in a cell culture incubator (37°C, 5% CO_2_). Before removing medium and washing cells with the above automated protocol, cell viability was measured using the RealTime-Glo™ MT Cell Viability Assay (Promega Cat. #G9713) following the manufacturer’s instructions. Briefly a 5X stock of reagent was prepared, 10 µl was added per well, and the plate was incubated at 37°C for 30 minutes before reading luminescence. Media was then removed, and the cells were washed using the automated protocol. Glycogen was measured using the automated protocol described above.

## ASSOCIATED CONTENT

### Supporting Information

Supporting Information at the end of the article includes: Glycogen detection assay performance (Figure S1); glycogen accumulation as a function of glucose concentration (Figure S2); validation of the automated protocol (Figure S3); layout of the inhibitor dilution and cell-based screening plates (Figure S4); performance of quality control compound C1 during screening (Figure S5); outline of automated screening protocol (Table S1)

## AUTHOR INFORMATION

### Author Contributions

Conceptualization: D.L., R.C., H.M., D.T.B., J.V.

Experiment Design: D.L., R.C., G.V., H.M., D.T.B., J.V.

Experimentation: D.L., R.C., G.V., H.M., K.T.M., J.V.

Data Analysis: D.L., G.V., J.V.

Writing-Original Draft and Editing: D.L., R.C., J.C.U., J.V.

Writing-Review: D.L., R.C., G.V., H.M., K.T.M., D.T.B., J.C.U., J.V.

## SUPPORTING INFORMATION

**Figure S1.**
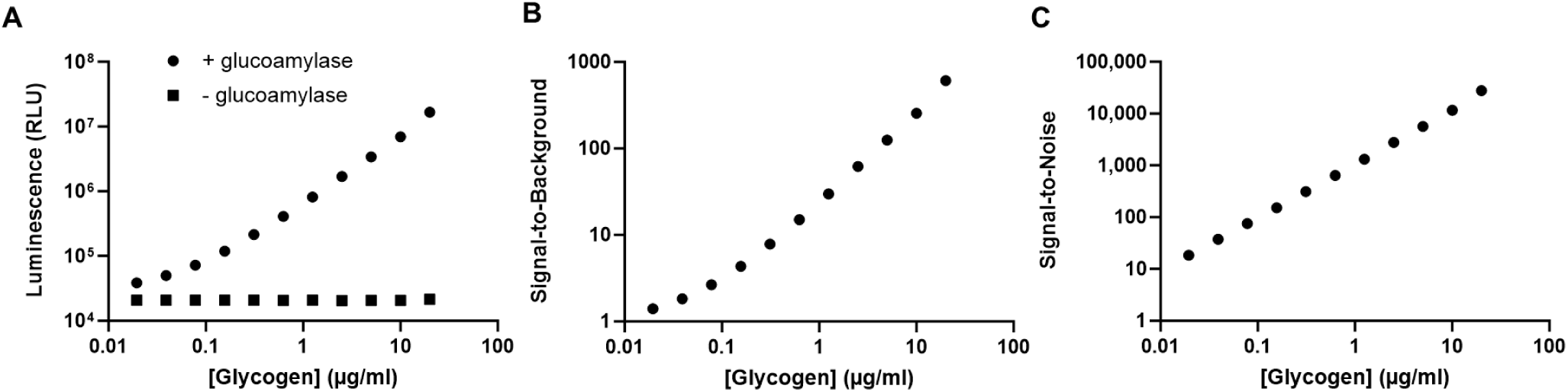
Parameters of the glycogen detection assay. (A) A purified stock of glycogen was serially diluted two-fold in PBS, starting from 20 µg/ml, and assayed in the presence and absence of glucoamylase. A negative control (assay background) containing PBS without glycogen was included. Each concentration was tested in quadruplicate in the wells of a 96-well plate. The average RLU are plotted. Error bars are +/− 1 s.d. Percent CVs were <5%. (B) The average RLU was used to calculate signal-to-background ratios. (C) The average net RLU and standard deviation of the background control were used to calculate the signal-to-noise ratio.

**Figure S2:**
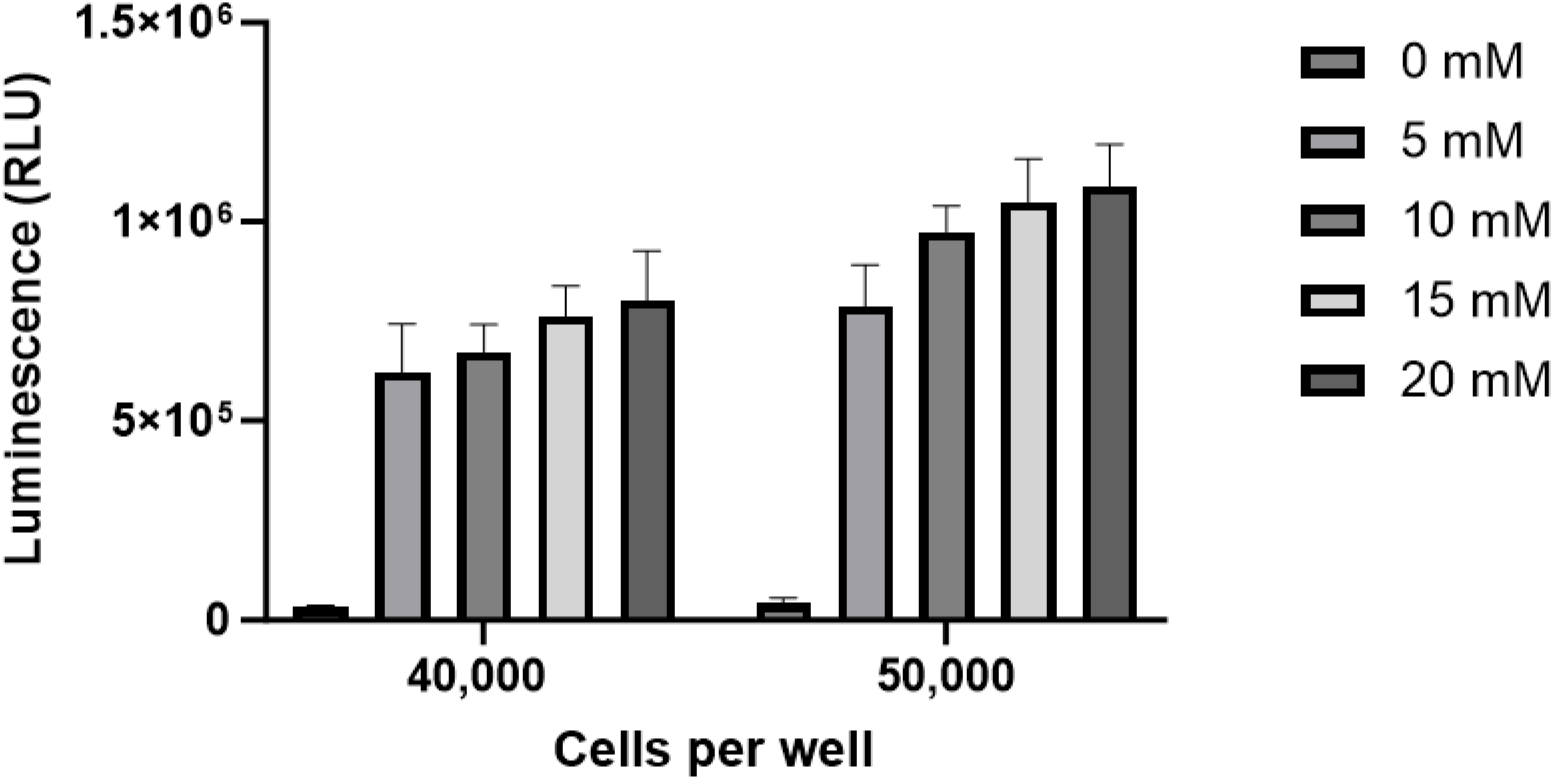
Accumulation of glycogen. After an overnight starvation in flasks, HeLa cells were collected and washed and plated in the wells of a 96-well plate at the indicated cell number per well. They were then incubated overnight in the presence of medium containing 0 to 20 mM glucose before measuring glycogen. Each condition was assayed in quadruplicate and the average RLU are plotted. Error bars are +/− 1 s.d.

**Figure S3.**
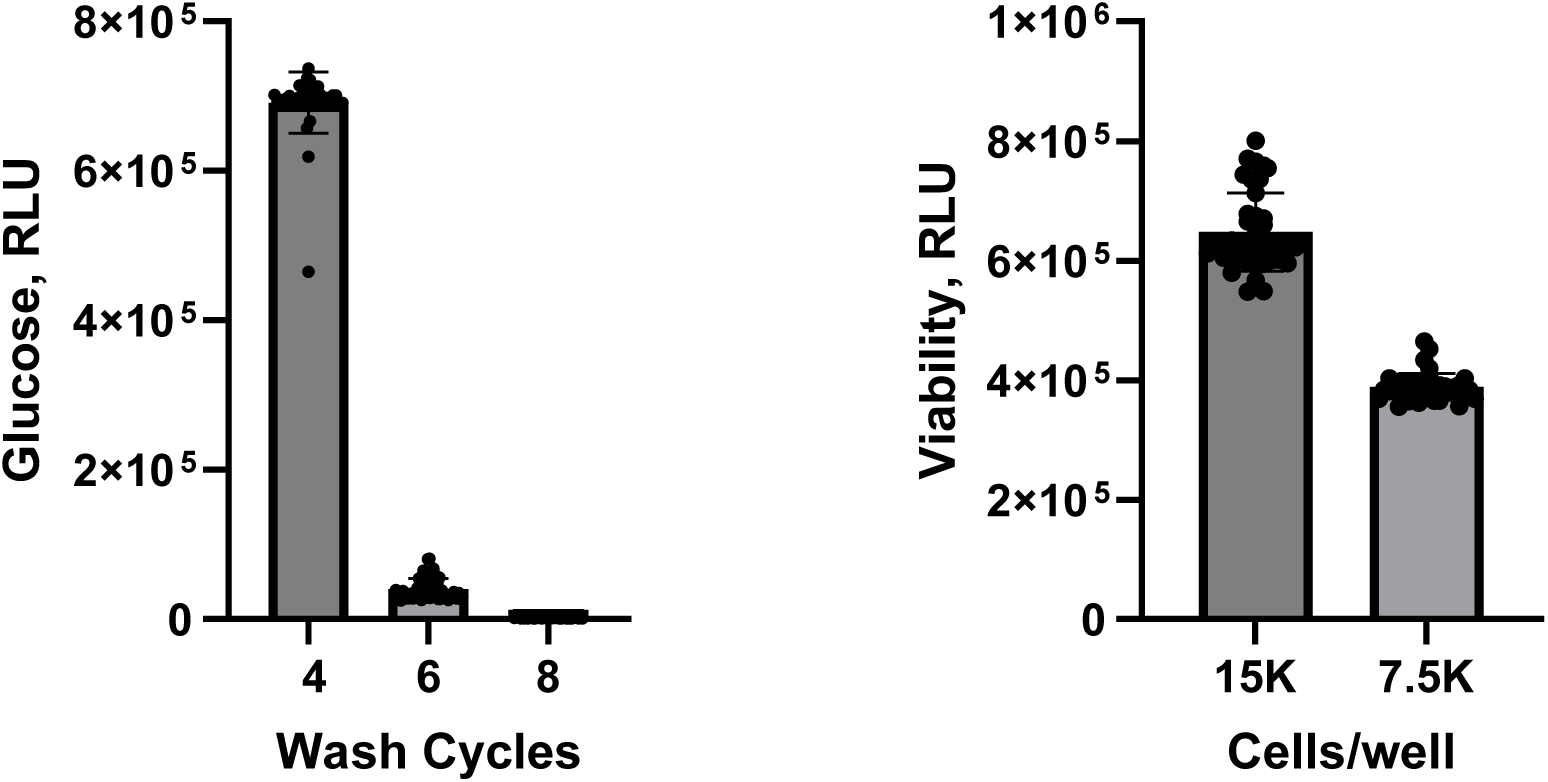
Validation of the automated protocol. (A) Measurement of residual glucose after the indicated number of wash cycles. (B) Viability measurements after the 8^th^ wash cycle using the CellTiter-Glo^®^ Assay. In both panels, each data set is for 40 wells.

**Figure S4.**
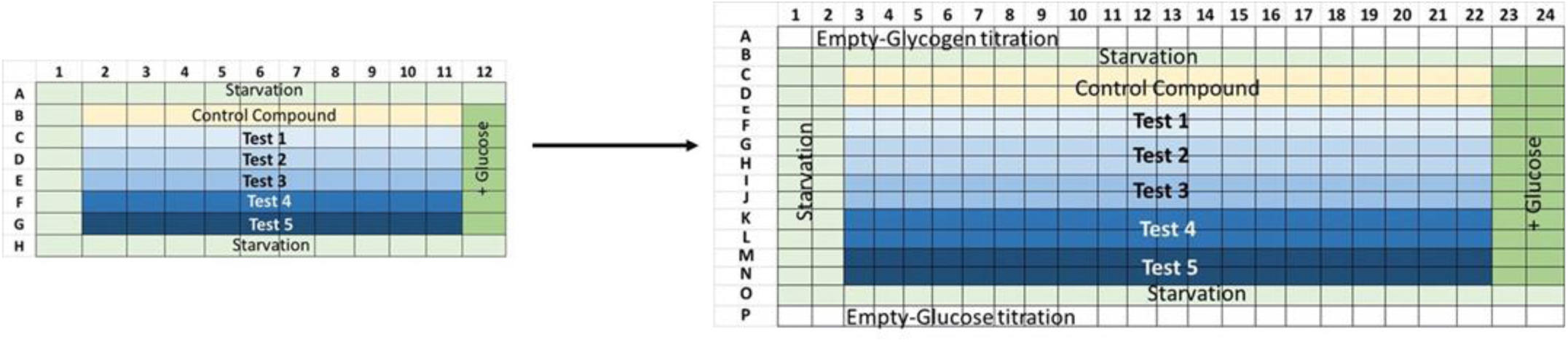
Layout of inhibitor dilution (96-well) and cell-based screening (384-well) plates. Compounds were diluted in each 96-well plate and then transferred to a corresponding 384-well plate. Rows A and P were reserved for glycogen and glucose titrations in the final assay plates.

**Figure S5.**
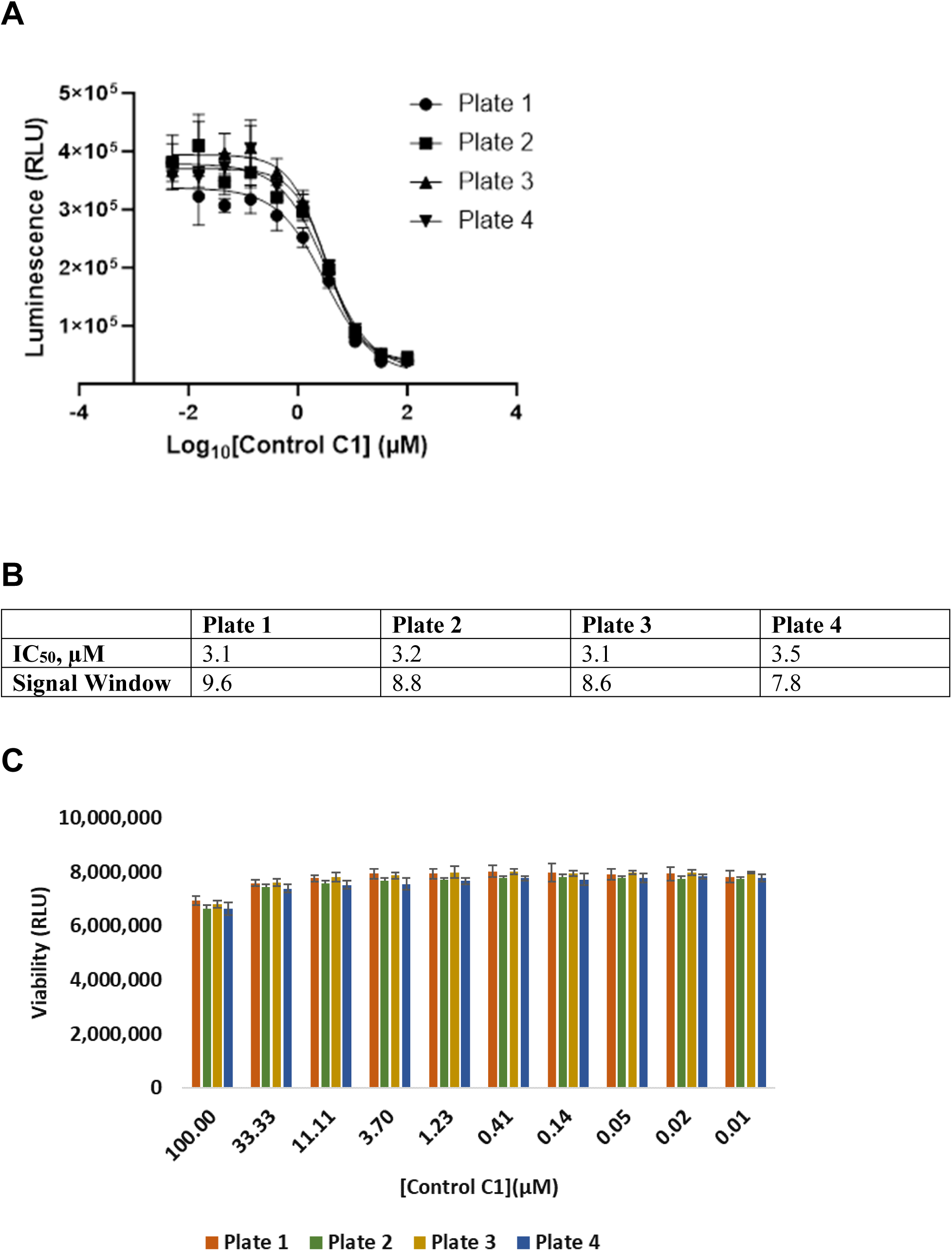
Control compound C1 performance across four plates. (A) Control compound C1 was assayed in a 10-point dose response curve on each of the four screening plates in Figure 6. Each concentration was tested in quadruplicate. The average RLU was calculated. Error bars are +/− 1 s.d. (B) The IC_50_ and maximum signal window for each of the plates. The signal window was calculated as the ratio of signal at low and high compound concentrations. (C) Cell viability was assessed using the RealTime-Glo™ Assay prior to cell lysis as described in the Methods section.

**Table S1:** Automated Screening Protocol.

|  | Step | Volume, time | Details |
| --- | --- | --- | --- |
| 1 | Cells | 24 h | Culture in T75 flasks overnight in starvation medium. |
| 2 | Cells | 25 µl | Collect cells, wash, and resuspend to 600,000 cells/ml in starvation medium.<br>Transfer 25 µl (15,000 cells) to the wells of a 384-well plate. |
| 3 | Library Compounds |  | 10mM compounds in DMSO.<br>Prepare 10-point dilution series starting with 100 µM (or 60 µM) compound in recovery medium with 20 mM glucose and 2% DMSO. |
| 4 | Add Compounds to Cells | 25 µl | Transfer 25 µl to cells in 384-well plate. (Cells are now in 10 mM glucose.)<br>Assay each compound dilution in quadruplicate wells. |
| 5 | Incubate | 24 h | 37°C, 5% CO<sub>2</sub> |
| 6 | Cell Viability measurement | 30-60 min | RealTime-Glo™ Assay<br>Bioluminescence |
| 7 | Wash | 8 cycles | Remove media.<br>Wash with PBS, 70 µl per wash cycle.<br>After final wash, remove PBS, leaving 15 µl. |
| 8 | Lyse | 10 min | Add 7.5 µl 0.3N HCl to cells.<br>Followed by addition of 7.5 µl 450 mM Tris pH 8.0. |
| 9 | Freeze plates | overnight | Seal plates and freeze overnight at -20°C |
| 10 | Glycogen digestion | 30 µl | Thaw cell lysate plates.<br>Prepare glucoamylase solution and add 30 µL to well containing cell lysate. |
| 11 | Incubation | 1 h | 37°C or Room temperature |
| 12 | Glucose measurement | 8 µl | Transfer 8 µl to wells of 384-well plate for the glucose detection assay. |
| 13 | Glycogen Assay readout | 90 min | Add 8 µl glucose detection reagent.<br>Incubate 90 min at RT.<br>Bioluminescence |

